# Polyketide synthase-derived sphingolipids determine microbiota-mediated protection against pathogens in *C. elegans*

**DOI:** 10.1101/2024.02.06.579051

**Authors:** Lena Peters, Moritz Drechsler, Barbara Pees, Georgia Angelidou, Liesa Salzer, Karlis Arturs Moors, Nicole Paczia, Hinrich Schulenburg, Yi-Ming Shi, Christoph Kaleta, Michael Witting, Helge B. Bode, Katja Dierking

## Abstract

Protection against pathogens is a major function of the gut microbiota. Although bacterial natural products have emerged as crucial components of host-microbiota interactions, their exact role in microbiota-mediated protection is largely unexplored. We addressed this knowledge gap with the nematode *Caenorhabditis elegans* and its microbiota isolate *Pseudomonas fluorescens* MYb115 that is known to protect against *Bacillus thuringiensis* (Bt) infection. We find that MYb115-mediated protection depends on sphingolipids that are derived from an iterative type I polyketide synthase (PKS), thereby describing a noncanonical pathway of bacterial sphingolipid production. We provide evidence that MYb115-derived sphingolipids affect *C. elegans* tolerance to Bt infection by altering host sphingolipid metabolism. This work establishes sphingolipids as structural outputs of bacterial PKS and highlights the role of microbiota-derived sphingolipids in host protection against pathogens.

## Introduction

A major function of the gut microbiota is its contribution to host protection against pathogens ^1^. The protective mechanisms conferred by the gut microbiota are complex and include direct competitive or antagonistic microbe–microbe interactions and indirect microbe-host interactions, which are mediated by the stimulation of the host immune response, promotion of mucus production, and maintenance of epithelial barrier integrity ^2^. Microbiota-derived metabolites are known to play an important role in the crosstalk between the gut microbiota and the immune system ^3–5^. Of these metabolites, bacterial natural products have emerged as crucial components of host-microbiota interactions ^6–8^.

Bacterial natural products (also called secondary or specialised metabolites) are chemically distinct, often bioactive compounds that are not required for viability, but mediate microbial and environmental interactions^9^. Some of the most studied natural products include polyketides (PKs), which are derived from polyketide synthase (PKS). PKS are found in many bacteria, fungi, and plants, and produce structurally diverse compounds by using an assembly line mechanism similar to fatty acid synthases ^10^. Many PKS-derived natural products show potent antibiotic (e.g., erythromycin and tetracycline), antifungal (e.g., amphotericin and griseofulvin) or immunosuppressant (e.g. rapamycin) activities ^11^ and have thus long played a central role in advancing therapeutic treatments for a wide range of medical conditions. The majority of characterised PKs were isolated from free-living microbes, while only a few are known to be gut microbiota-derived ^8^. Most well-studied examples of PKS-derived products from the microbiota are virulence factors associated with pathogenicity ^12^. Few PKS-encoded natural products were reported to play a role in microbiota-mediated protection against pathogens both directly and indirectly. For example, the antifungal PK lagriamide supports direct symbiont-mediated defence of eggs against fungal infection in the beetle *Lagria vilossa* ^13^. A PKS cluster of the rodent gut symbiont *Limosilactobacillus reuteri* is required for activating the mammalian aryl hydrocarbon receptor (AhR), which is involved in mucosal immunity ^14^. Additionally, *L. reuteri* PKS was recently demonstrated to exhibit antimicrobial activity and to drive interspecies antagonism ^15^. Yet, the vast majority of microbiota encoded PKS are of unknown function and mechanistic studies linking specific microbial natural products to host phenotypes are scarce.

The *Pseudomonas fluorescens* isolate MYb115 belongs to the natural gut microbiota of the model organism *Caenorhabditis elegans* ^16^. It was previously found that MYb115 protects *C. elegans* against the harmful effects of infection with *Bacillus thuringiensis* (Bt) without directly inhibiting pathogen growth, likely through an indirect, host-dependent mechanism ^17,18^. The nature of the microbiota-derived protective molecule and the involved host processes were unknown. Here, we identify a biosynthetic gene cluster (BGC) in MYb115 encoding an iterative type I polyketide synthase (PKS) that is required for MYb115-mediated protection and produces sphingolipids. We demonstrate that MYb115-derived sphingolipids affect *C. elegans* sphingolipid metabolism and establish the importance of *C. elegans* sphingolipid metabolism for survival after Bt infection.

## Results

### *P. fluorescens* MYb115 PKS is required for *C. elegans* protection against Bt infection

The natural microbiota isolate *P. fluorescens* MYb115 protects *C. elegans* against infection with the Gram-positive pathogenic *B. thuringiensis* strain Bt247 likely through a host-dependent mechanism ^18^, but the nature of the microbiota-derived protective molecule was unknown. We performed an antiSMASH analysis^19^ of the MYb115 genome to identify natural product biosynthetic gene clusters (BGCs). We found three BGCs in the MYb115 genome, encoding a non-ribosomal peptide synthetase (NRPS), an iterative type I polyketide synthase (PKS) cluster, and an arylpolyene pathway.

We modified the PKS and NRPS clusters of MYb115 by inserting the inducible arabinose P*_BAD_* promoter. Thus, while induction of BGC expression requires arabinose supplementation, no expression is observed in the absence of arabinose supplementation, mimicking a deletion phenotype ^20^. We assessed the ability of MYb115 P*_BAD_sga* (MYb115 PKS cluster) and MYb115 P*_BAD_nrpA* (MYb115 NRPS cluster) in an induced (+ arabinose) and non-induced (-arabinose) state to protect *C. elegans* against Bt247 infection. We found that infected *C. elegans* exposed to induced MYb115 P*_BAD_sga* showed significantly increased survival when compared to infected worms on MYb115 P*_BAD_sga* in a non-induced state (Figure 1A, Table S1). Supplementation of the *C. elegans* laboratory food *Escherichia coli* OP50 with arabinose did not affect survival, showing that arabinose itself does not influence *C. elegans* resistance to Bt (Table S1). While the PKS gene cluster affects MYb115-mediated protection, we did not observe significant differences in worm survival with or without arabinose supplementation on the MYb115 P*_BAD_nrpA* strain (Figure 1B, Table S1), indicating that the MYb115 NRPS gene cluster is not involved in MYb115-mediated protection against Bt infection.

**Figure 1:**
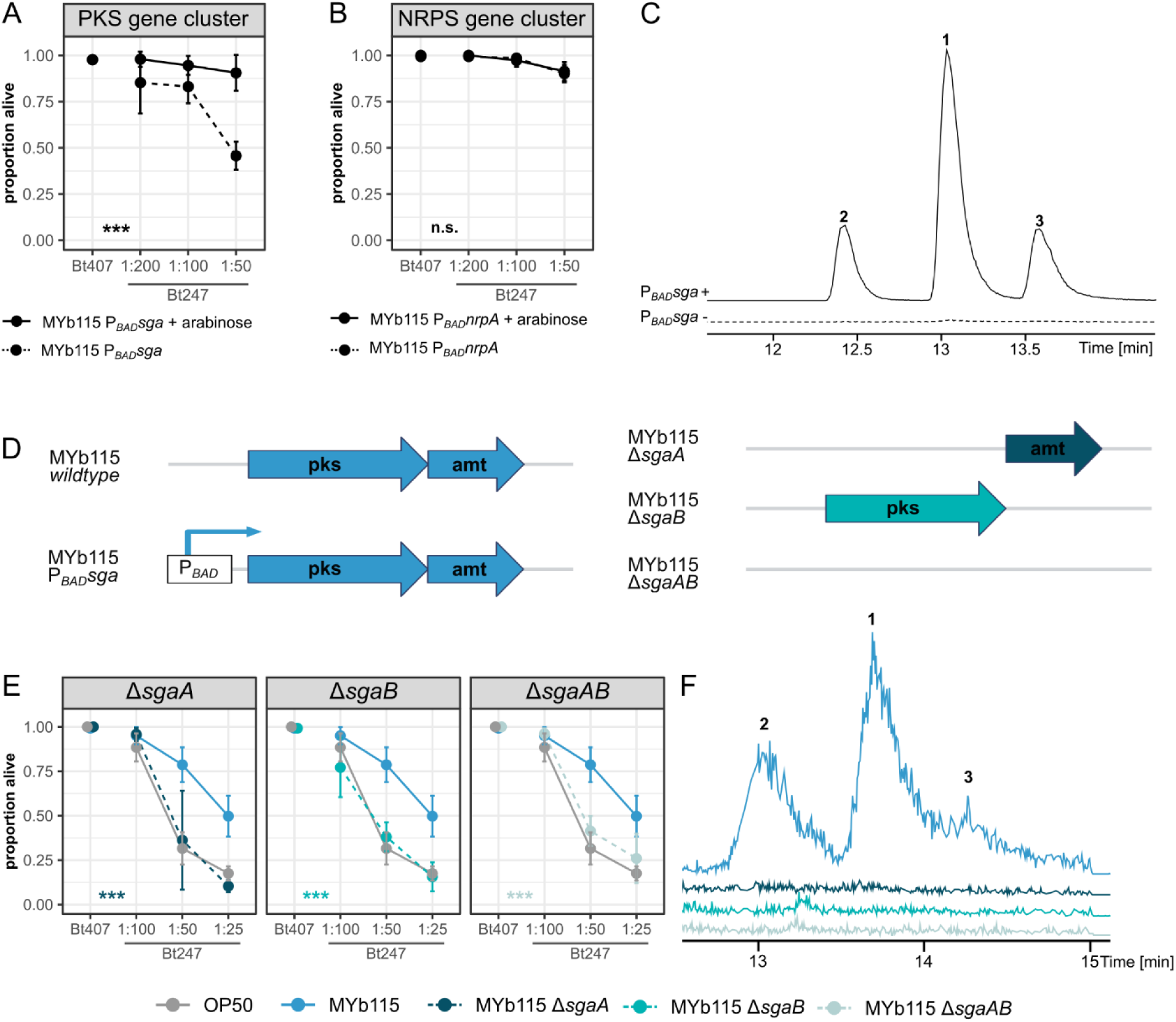
MYb115 PKS-derived sphingolipids mediate protection against *B. thuringiensis* infection. **(A, B)** Survival proportion of *C. elegans* N2 on *P. fluorescens* MYb115 P*BADsga* **(A)** or MYb115 P*BADnrpA* **(B)** induced with arabinose (solid line) or in a non-induced state without arabinose supplementation (dashed line) 24 h post infection with *B*. *thuringiensis* Bt247. Bt407 was used as a non-pathogenic control. The data shown is representative of three independent runs with four replicates each (see Table S1). **(C)** LC-MS chromatogram of MYb115 P*BADsga* extracts from cultures with (solid line) and without (dashed line) arabinose supplementation. Upon induction with arabinose, three compounds (**1-3**) are produced. **(D)** Schematic representation of the MYb115 PKS gene cluster and its modifications. Polyketide synthase (pks) SgaA, aminotransferase (amt) SgaB, inducible arabinose promoter (P*BAD*). **(E)** Survival proportion of N2 on *E. coli* OP50, MYb115, or MYb115 knockout mutants. *C. elegans* on all three tested mutants MYb115 Δ*sgaAB*, MYb115 Δ*sgaA*, and MYb115 Δ*sgaB* were significantly more susceptible to infection with Bt247 than worms on wildtype MYb115. Means ± standard deviation (SD) of n = 4, are shown in all survival assays **(A, B, E)**. Statistical analyses were carried out with the GLM framework and Bonferroni adjustment for multiple testing, \*\*\**p* < 0.001. **(F)** LC-MS chromatogram of MYb115 wt, Δ*sgaA*, Δ*sgaB* and Δ*sgaAB*. Production of compounds **1-3** was abolished in all three deletion mutants. Raw counts and additional survival runs can be found in Table S1.

We then deleted either the entire gene cluster (MYb115 Δ*sgaAB*), the polyketide synthase SgaA (MYb115 Δ*sgaA)*, or the aminotransferase SgaB (MYb115 Δ*sgaB*) (Figure 1D) to confirm the requirement of the MYb115 PKS cluster in MYb115-mediated protection. While MYb115 provided significant protection against infection in *C. elegans* compared to worms on *E. coli* OP50 (Figure 1E, ^18^), protection of worms on all three MYb115 mutants was lost (Figure 1E).

### *P. fluorescens* MYb115 PKS produces long chain sphinganines and phosphoglycerol sphingolipids

MYb115-mediated protection against Bt247 infection depends on the MYb115 PKS SgaAB. We next asked which natural product is produced by SgaAB. Using LC-MS, we identified three compounds that are produced in MYb115 P*_BAD_sga* upon induction with arabinose (Figure 1C). We subsequently established that the compounds **1**-**3** are also produced, but less abundant, in MYb115, and that all three MYb115 deletion mutants (MYb115 Δ*sgaAB*, MYb115 Δ*sgaA*, and MYb115 Δ*sgaB*) are not able to produce compounds **1**-**3** (Figure 1F). MS^2^ experiments revealed that compounds **1**-**3** show structural similarities to commercially available long chain sphinganines (Figure S1). We determined the molecular composition through isotopic labeling experiments and confirmed that compounds **1**-**3** are very long chain sphinganines (Figure S2A-C).

To test if the MYb115-derived sphinganines exist as free compounds or are part of lipids, we performed lipidomic analysis of MYb115. We found that in addition to the three sphinganines **1**-**3** MYb115 produces compounds **4**-**6**, each with masses 154 Da heavier than those of the three sphinganine derivatives (Figure 2A, S2D-F). Since the masses of **4**-**6** did not match any known lipids in the MS-DIAL LipidBlast (version 68) dataset, we used the exact mass and different lipid headgroups to propose structures for compounds **4**-**6**. We conclude that compounds **4**-**6** are most likely phosphoglycerol sphingolipids (PG-sphingolipids) (Figure 2A). Next, we analysed the relative abundance of sphinganines **1**-**3**, and PG-sphingolipids **4**-**6** in MYb115 and MYb115 P*_BAD_sga* induced by arabinose or repressed by glucose supplementation (Figure S2G). While the sphinganines **1**-**3** were more abundant in the induced MYb115 P*_BAD_sga* samples, the total abundance of PG-sphingolipids **4** and **5** did not differ compared to MYb115 supplemented with arabinose (Figure S2G). Thus, increase in sphinganine production does not necessarily lead to increase in PG-sphingolipid production. However, since the MYb115 deletion mutants MYb115 Δ*sgaAB*, MYb115 Δ*sgaA*, and MYb115 Δ*sgaB* do not produce sphinganines, and thus likely also not PG-sphingolipids, we are not able to differentiate between the effects of the individual sphingolipid species.

**Figure 2:**
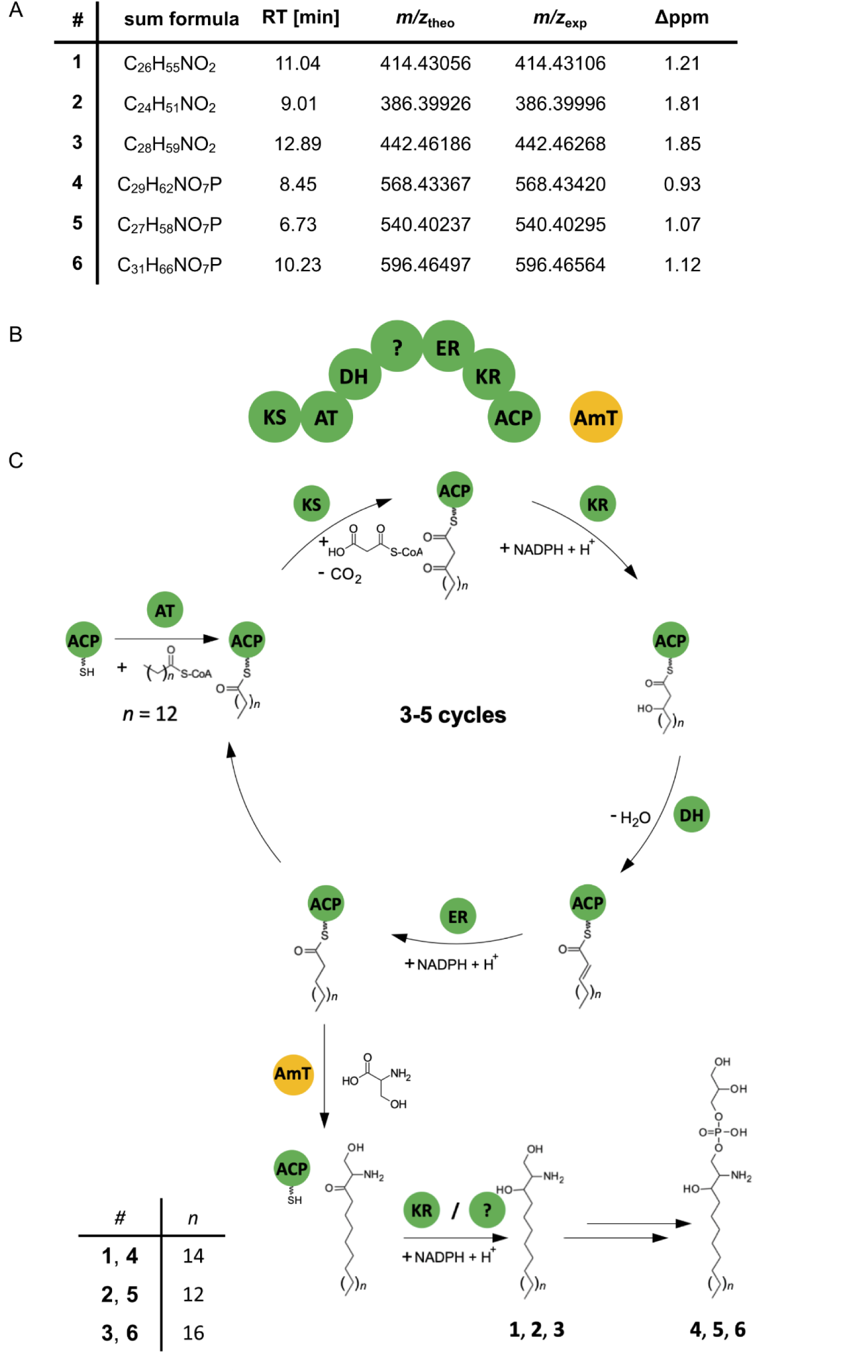
Structure and proposed biosynthesis of *P. fluorescens* MYb115 PKS SgaAB-derived sphingolipids. **(A)** Sum formula and LC-MS/MS data of MYb115-derived sphinganines **1**-**3** and phosphoglycerol sphingolipids **4**-**6** discovered by lipidomic analysis of MYb115. Retention time in minutes (RT [min]); mass-to-charge ration (m/z); theoretical mass (theo), experimental mass (exp); mass error in parts per million (Δppm). **(B)** Schematic representation of the PKS SgaA domains (green) and the aminotransferase SgaB (yellow). **(C)** Possible biosynthesis scheme of long chain sphinganines **1**, **2** and **3** and the phosphoglycerol sphingolipids **4**, **5** and **6**. First, a palmitoyl-CoA starter unit is extended and reduced in 3-5 cycles. The resulting ACP domain-bound fatty acid is subsequently connected to serine in a reaction catalyzed by SgaB. The final remaining carbonyl group is reduced by the ketoreductase or the cryptic domain. Finally, the resulting long chain sphinganine is bound to a phosphoglycerol-head group. ACP = acyl-carrier protein; AmT = aminotransferase; AT = acyl transferase; DH = dehydratase; ER = enoyl reductase; KR = ketoreductase; KS = ketosynthase;= cryptic domain. Only the domains responsible for the respective reactions are shown.

### A proposed pathway for iT1PKS-dependent sphingolipid biosynthesis

Sphingolipid synthesis in bacteria and eukaryotes involves the condensation of an amino acid (typically serine in mammals) and a fatty acid (typically palmitate in mammals) *via* the serine palmitoyl transferase (SPT) enzyme that uses pyridoxal phosphate as cofactor for serine decarboxylation and coupling to palmitoyl-CoA ^21^. In the case of MYb115, which lacks the SPT gene, the protective sphingolipids are produced by the two-gene cluster *sgaAB* (**S**phin**ga**nine biosynthesis **A** and **B**), in which *sgaA* encodes an PKS and *sgaB* encodes a pyridoxal-dependent protein with similarity to 2-amino-3-ketobutyrate CoA ligase (KBL) or aminotransferase (AMT) (Figure 2B), most likely substituting the SPT function. The full reductive loop of SgaA suggests elongation of palmitoyl-CoA by 3-5 cycles of subsequent polyketide elongation and reduction with malonyl-CoA as extender unit leading to an acyl carrier protein (ACP) domain-bound C22-C26 fatty acid still bound to SgaA, which is then connected to serine by SgaB in a pyridoxal phosphate dependent and SPT-like manner (Figure 2C). Through isotopic labelling experiments we could show that ^13^C^15^N-labelled serine is indeed incorporated during sphinganine biosynthesis in MYb115 (Table S2).

### Homologous PKS are present across diverse bacterial genera

Iterative PKS were originally found in fungi and only rarely in bacteria ^10^. However, a large number of bacterial iterative PKS were identified more recently ^22^. While only a few bacterial iterative PKS and their products have been studied, our work is to our knowledge the first example of a PKS shown to be involved in sphingolipid biosynthesis and also the first description of a *P. fluorescens* isolate as sphingolipid producer. We explored the distribution of the two-gene MYb115 PKS SgaAB in bacteria and found 6,101 homologous putative PKS (Table S3). Interestingly, the homologous PKS were present in bacteria that are known to be closely associated with hosts, including human pathogens and opportunistic pathogens (Figure 3A). When we analysed the distribution of the target BGC class at the genus level, we found that the putative PKS is dominantly distributed in *Burkholderia* (Figure 3B).

**Figure 3:**
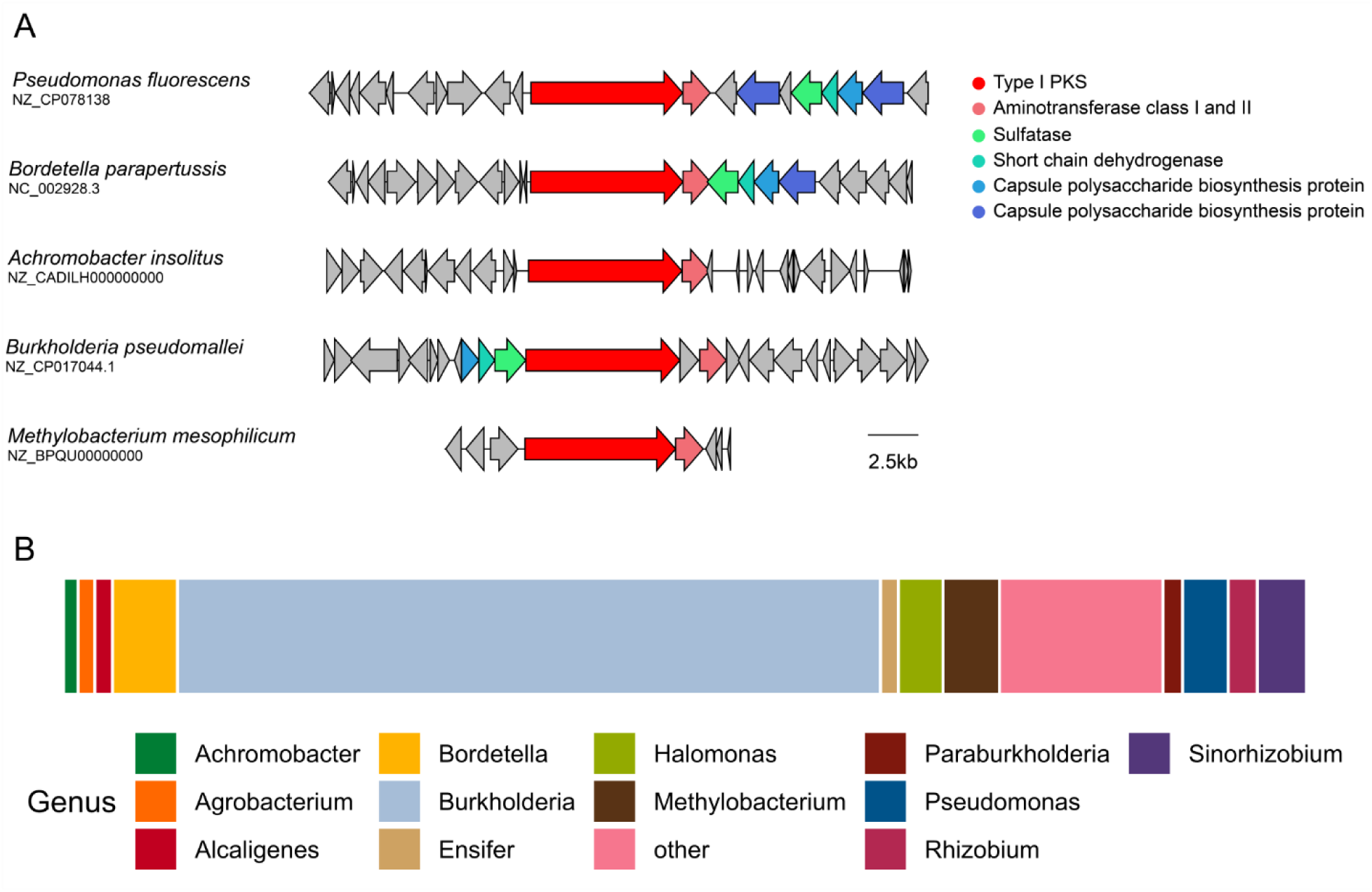
Distribution of *P. fluorescens* MYb115 PKS SgaAB homologues in bacteria. The monomodular PKS (KW062_RS19805) and the aminotransferase (KW062_RS19800) in *P. fluorescens* MYb115 (NZ_CP078138) were searched against the NR NCBI database (https://www.ncbi.nlm.nih.gov/) using cblaster (1.8.1) ^23^. **(A)** Five representative PKS SgaAB homologs from various bacterial genera aligned and visualised using clinker ^24^. **(B)** Total distribution of 6,101 PKS SgaAB homologs across different bacterial genera.

### MYb115-derived sphingolipids alter host fatty acid and sphingolipid metabolism

In a first step towards exploring the function of microbiota-derived sphingolipids in mediating the interaction with the host, we tested whether MYb115-produced sphingolipids affect the ability of MYb115 to colonize the host or modulate host feeding behavior. We did not observe a difference in host colonization between MYb115 and Δ*sgaAB* MYb115 (Figure S3A, Table S4), nor did we see differences in *C. elegans* feeding behavior on MYb115 and Δ*sgaAB* MYb115 (Figure S3B, Table S4).

Mouse lipid metabolism was previously shown to be affected by gut microbiota-derived sphingolipids ^25^. Moreover, in a *C. elegans* Parkinson disease model, the probiotic *B. subtilis* strain PXN21 protects the host against protein aggregation by modulating sphingolipid metabolism ^26^. Thus, we hypothesied that MYb115-derived sphingolipids impact host metabolism. To test this hypothesis, we performed gene expression profiling of 1-day adult worms on either MYb115 or MYb115 Δ*sgaAB* in the absence and presence of pathogenic Bt247. We did not observe any genes differentially regulated between worms on sphingolipid-producing MYb115 and worms on the MYb115 Δ*sgaAB* mutant when using an adjusted *p*-value cutoff of 0.05. However, integrating the transcriptomic data into the iCEL1314 genome-scale metabolic model of *C*.

*elegans* ^27^ to create context-specific models, resulted in 24 and 23 significant differences in the presence or absence of Bt247, respectively. Through a pathway enrichment analysis, we found that in the absence of Bt247, animals colonized by MYb115 or MYb115 Δ*sgaAB* varied in the activity of multiple pathways linked with sphingolipid precursor production, such as fatty acid biosynthesis and elongation, as well as sphingolipid metabolism itself (Figure 4A, Table S5). In the presence of Bt247, colonization with either MYb115 or MYb115 Δ*sgaAB* affected most strongly propanoate metabolism (Figure S4A). Here, we also saw an enrichment in valine, leucine, and isoleucine degradation, which are branched-chain amino acids (BCAA). This pathway is directly connected with propanoate metabolism that provides components for the synthesis of the C15iso fatty acid, which is the precursor for sphingolipids in *C. elegans* ^28^. Focusing on sphingolipid metabolism, a flux variability analysis ^29^ revealed a significant difference in upper bound values for the sphingolipid metabolism reactions in worms infected with Bt247 on MYb115 *versus* MYb115 Δ*sgaAB* (t-test *p*-value=0.00033). Among those reactions, six reactions that all have ceramide as a substrate or product had the strongest changes (Figure S4B). Overall, these findings suggest that worms colonized by MYb115 *versus* MYb115 Δ*sgaAB* have a significantly reduced capacity to generate sphingolipids.

**Figure 4:**
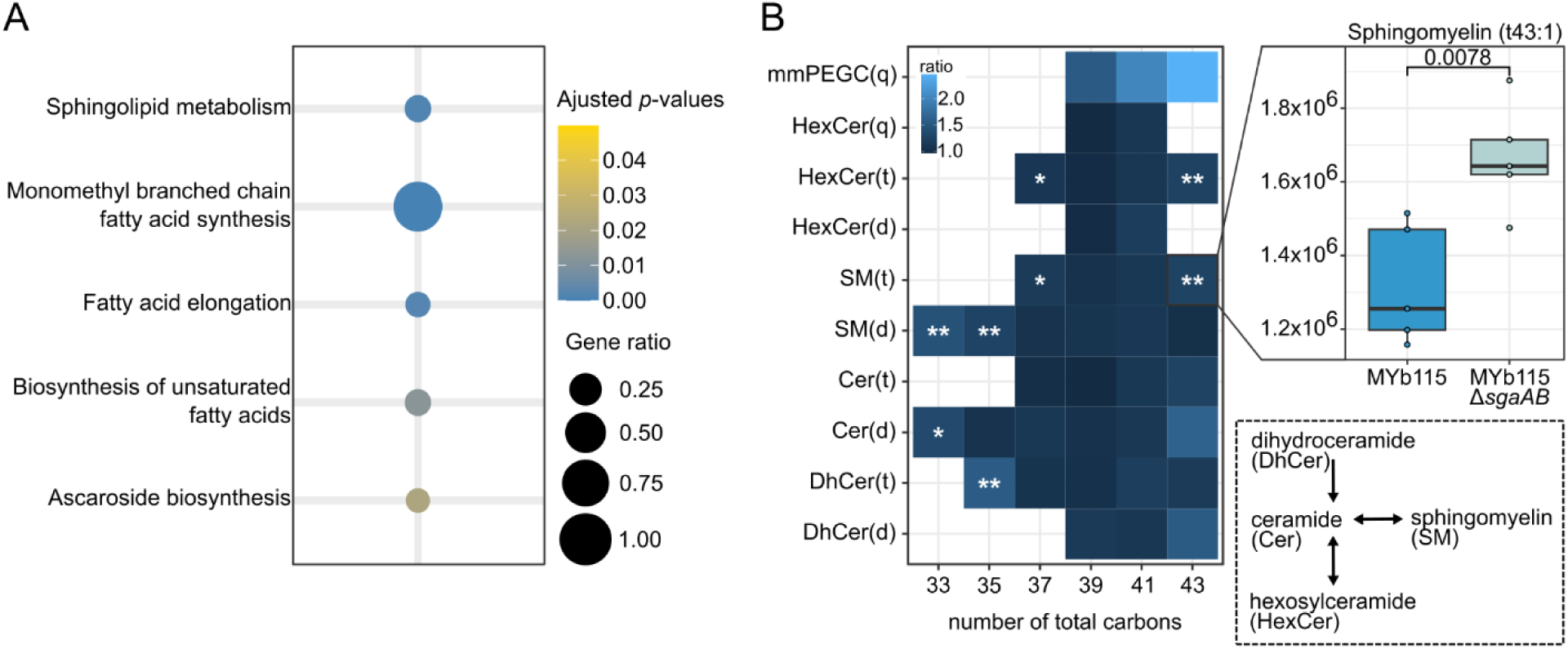
MYb115-derived sphingolipids modulate host sphingolipid metabolism. **(A)** Enriched metabolic subsystems identified by Flux enrichment analysis based on transcriptome data comparing worms treated with MYb115 and MYb115 Δ*sgaAB*. Significant reactions from linear regression models of all 3 data types (upper bound, lower bound, Optimal Flux Distribution) were combined (while removing duplicates) and used against the background of all reactions within the iCEL1314 *C. elegans* metabolic model. Enrichment was performed with the FEA function in the COBRA toolbox. **(B)** Reduced sphingolipid contents in worms exposed to MYb115 compared to worms exposed to MYb115 Δ*sgaAB.* The heatmap shows the differences in ratio of detected sphingolipids between the mean of MYb115 Δ*sgaAB* and the mean of MYb115. The boxplot shows the difference in ratio of Sphingomyelin (t43:1) in worms exposed to MYb115 Δ*sgaAB* and MYb115, all remaining boxplots can be found in Figure S5. Statistical analysis was done with a Welch’s *t*-test, * *p*-value < 0.05, ** *p*-value < 0.01. Dihydroceramides (DhCer), Ceramides (Cer), Sphingomyelins (SM), Hexosylceramides (HexCer), with hydroxylated fatty acyls (t) or non-hydroxylated fatty acyls (d), Hexosylceramides with phytosphingosine base and hydroxylated fatty acyls (HexCer(q)), monomethyl phosphoethanolamine glucosylceramide (mmPEGC(q)).

### MYb115-derived sphingolipids interfere with *C. elegans* complex sphingolipids

The metabolic network analysis revealed that sphingolipid metabolism reactions show differential activity between MYb115 and MYb115 Δ*sgaAB*. To confirm that MYb115-derived sphingolipids affect *C. elegans* sphingolipid metabolism, we performed lipidomic profiling of *C. elegans* exposed to MYb115 or MYb115 Δ*sgaAB*. We identified *C. elegans* sphingolipids by manual interpretation of MS^1^ and MS^2^ data and used sphingolipids that have previously been described in *C. elegans* containing a C17iso-branched chain sphingoid base and different length of N-Acyl chains as input ^30^ (Table S6). Since the employed analytical method cannot separate between different hexoses attached to the sphingolipid they were annotated as hexosylceramides (HexCers), which showed the neutral loss of 162.052275 Da. Monomethylated phosphoethanolamine glucosylceramides (mmPEGCs), a class of *C. elegans* phosphorylated glycosphingolipids, were identified based on fragments as previously described ^31^.

We were not able to detect MYb115-derived sphinganine in worms on MYb115. Likewise, we did not detect any sphingolipids based on sphinganines produced by MYb115. A possible explanation is that bacterial sphinganine concentrations in worms are below the detection limit. However, we found different complex host sphingolipids based on the C17iso-branchend chain sphingoid base typical for *C. elegans* with N-acyl sides of length 16-26 without or with hydroxylation. In addition to previously established sphingolipids, we identified HexCer with an additional hydroxyl group instead of the double bond in the sphingoid base. In total, we identified 40 *C. elegans* sphingolipids from different sphingolipid classes. We did not observe a difference in *C. elegans* C17iso sphinganine or C17iso sphingosine, but in certain dihydroceramide (DhCer) and ceramide (Cer) species between worms on MYb115 or MYb115 Δ*sgaAB*. Also, complex sphingolipids downstream of ceramides, i.e., sphingomyelins (SMs) and HexCers were increased in worms on MYb115 Δ*sgaAB*, and some even significantly increased (Figure 4B). All changes in *C. elegans* sphingolipids between worms on MYb115 and MYb115 Δ*sgaAB* are summarised in Figure 4B. Individual sphingolipid profiles are shown in Figure S5. Most of the significant changes occurred at the lower or upper end of the detected N-acyl chain length. No changes occurred in sphingolipids containing an N-acyl of 22 or 24 carbon length. However, the series of SM(d33:1, d35:1, d37:1), showed a consistent and significant increase. Additionally, SM(t37:1) and SM(t43:1) as well as the corresponding HexCer(t37:1) and SM(t37:1) increased significantly. Notably, we found the highest fold-changes between MYb115 and MYb115 Δ*sgaAB*-exposed worms for mmPEGC. However, changes were not significant and so far, the biosynthesis pathway of mmPEGCs is unknown.

Together, our data suggest that MYb115-derived sphingolipids interfere with *C. elegans* sphingolipid metabolism mainly at the conversion of dihydroceramide and ceramide to sphingomyelins and hexosylceramides.

### Modifications in *C. elegans* sphingolipid metabolism affect defence against Bt247 infection

Since MYb115 affects host sphingolipid metabolism and protects the worm against Bt infection, we next asked whether alterations in nematode sphingolipid metabolism affect *C. elegans* survival following Bt infection. We performed survival experiments using several *C. elegans* mutants of sphingolipid metabolism enzymes (Figure 5A, Figure S6, Table S7). We assessed the general involvement of sphingolipid metabolism in the response to Bt infection in the presence of the non-protective lab food *E. coli* OP50. We found that mutants of the *C. elegans* serine palmitoyl transferases *sptl-1(ok1693)* and *sptl-3(ok1927)*, which catalyze the *de novo* synthesis of the C17iso sphingoid base, showed increased survival on Bt in comparison to wildtype N2 worms (Figure 5C). Also, ceramide synthase mutants *hyl-1(ok976)* and *hyl-2(ok1766)*) showed improved survival on Bt (Figure 5C). Moreover, the two ceramide metabolic gene mutants, namely *cgt-1(ok1045)* and *cerk-1(ok1252)* were more resistant to Bt infection (Figure 5C). *cgt-1* encodes one of three *C. elegans* ceramide glucosyltransferases that generate glucosylceramides (GlcCers). *cerk-1* is a predicted ceramide kinase that catalyzes the phosphorylation of ceramide to form ceramide-1-phosphate (C1P). In contrast, the *sms-1(ok2399)* mutant was more susceptible to Bt247 infection than wildtype worms. *sms-1* encodes a *C. elegans* sphingomyelin synthase that catalyzes the synthesis of sphingomyelin from ceramide. Accordingly, the *asm-3(ok1744)* mutant, which lacks the enzyme that breaks down sphingomyelin to ceramide, showed increased resistance to Bt247 (Figure 5C). Notably, the ceramidase mutants *asah-1(tm495)* and *asah-2(tm609)* were also significantly more susceptible to Bt247 infection than the *C. elegans* control (Figure 5C). *asah-1* encodes a *C. elegans* acid ceramidase that converts ceramide to C17iso-sphingosine, which is subsequently phosphorylated by the sphingosine kinase SPHK-1 to C17iso-sphingosine-1-phosphate ^32^. Together, these results suggest that inhibition of *de novo* synthesis of ceramide and inhibition of the conversion of ceramide to GlcCer or C1P increases survival of *C. elegans* infected with Bt247, while inhibition of the conversion of ceramide to sphingomyelin or sphingosine decreases survival of Bt247-infected animals.

**Figure 5:**
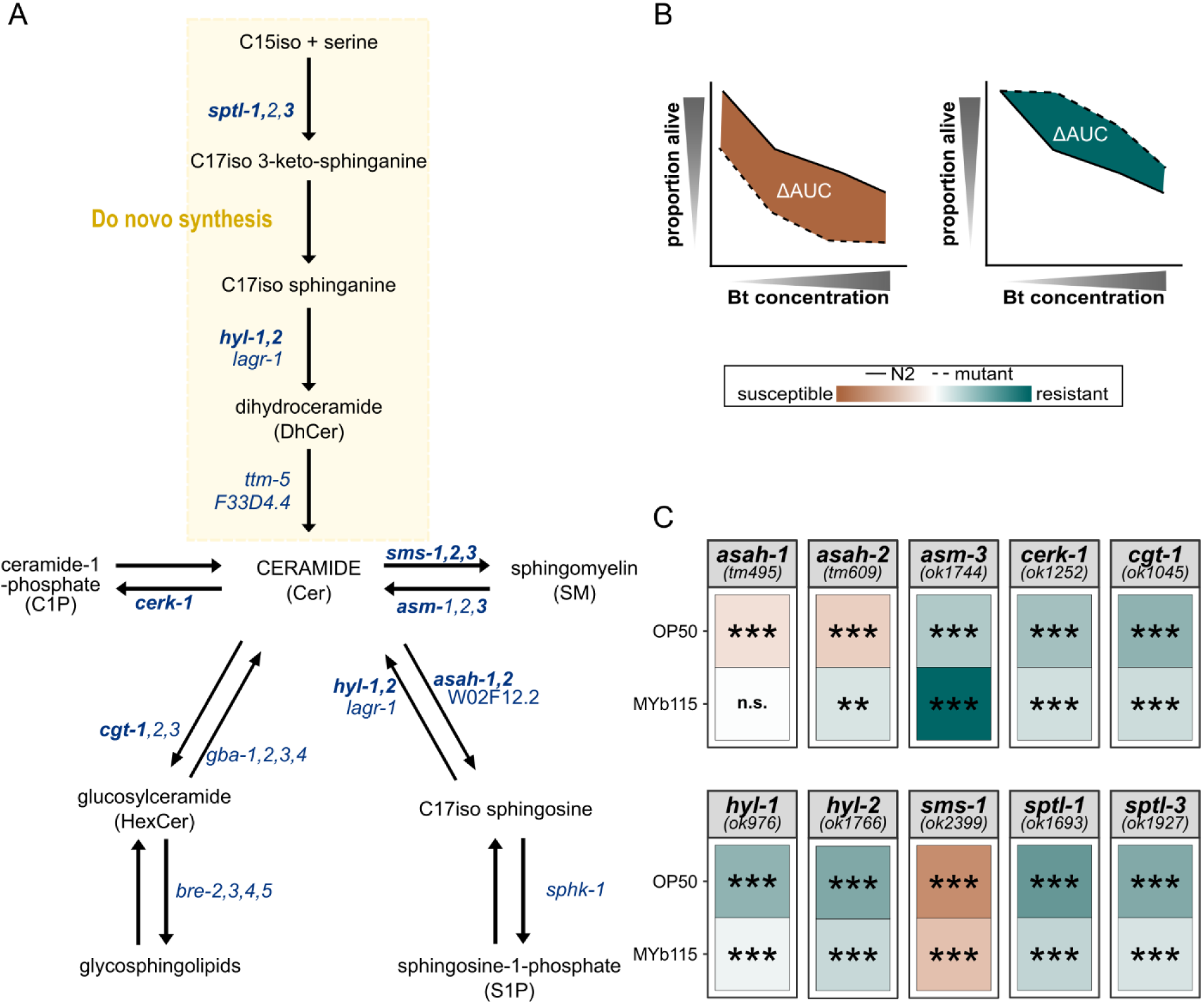
Modulations in *C. elegans* sphingolipid metabolism affect survival after Bt247 infection. **(A)** Overview of sphingolipid metabolism in *C. elegans*. *C. elegans* produces sphingoid bases which are derived from a C17 iso-branched fatty acid and are thus structurally distinct from those of other animals with mainly straight-chain C18 bases ^28^. *C. elegans* sphingolipids consist of a sphingoid base backbone derived from C15iso-CoA and serine, which is N-acylated with fatty acids of different lengths as well as different functional groups at the terminal hydroxyl group. Dihydroceramides (DhCers) are formed from C17iso sphinganine and fatty acids or 2-hydroxy fatty acids. Desaturation at the 4th carbon yields ceramides (Cers), which are the precursors of complex sphingolipids such as sphingomyelin (SM) and glucosylceramide (HexCer). Mutants of sphingolipid metabolism genes in bold were tested in survival assays shown in **(C)**. **(B)** Schematic survival comparing N2 wildtype (solid line) *versus* mutant strains (dashed lines), the difference of the area under the survival curve (AUC) is shaded in brown when the mutants are more susceptible to the infection than the control and in green when the mutants are more resistant to the infection. **(C)** Heatmap represents the ΔAUC of the survival of the *C. elegans* sphingolipid metabolism mutants *versus* average of the wildtype N2 strain. Here we summarised all conducted survival assays of either *E. coli* OP50- or *P. fluorescens* MYb115-treated worms infected with Bt247. Statistical analyses were carried out with the GLM framework and FDR adjustment for multiple testing, \*\*\**p* < 0.001. Each individual survival curve can be found in Figure S6A + B. Raw counts can be found in Table S7.

We additionally assessed *C. elegans* sphingolipid metabolism mutant survival on the protective microbiota isolate MYb115. MYb115 and the inhibition of *de novo* synthesis of ceramide or inhibition of the conversion of ceramide to GlcCer or C1P protect worms against infection with Bt247. Therefore, we did not expect to see an effect of MYb115 on the increased survival phenotype of the *sptl-1, -3*, *hyl-1, -2*, *cerk-1*, and *cgt-1* mutants. Our results are fully consistent with these expectations (Figure 5C). However, both ceramidase mutants *asah-1(tm495)* and *asah-2(tm609)*, which were more susceptible to Bt247 infection on *E. coli* OP50, were as susceptible as and even more resistant than wildtype worms on MYb115, respectively (Figure 5C). Notably, MYb115 also ameliorated the susceptibility phenotype of the *sms-1(ok2399)* mutant (Figure 5C). These data indicate that MYb115 interacts with host sphingolipid metabolism at least at the conversion of ceramide to sphingomyelin and C17iso-sphingosine.

## Discussion

Understanding microbiota-host interactions at the level of the molecular mechanism requires the identification of individual microbiota-derived molecules and their associated biological activities that mediate the interaction. In this study we demonstrate that *P. fluorescens* MYb115-mediated host protection ^18^, depends on bacterial-derived sphingolipids. We show that MYb115 produces protective sphingolipids by a biosynthetic gene cluster encoding an iterative PKS. This finding is important since eukaryotes and all currently known sphingolipid-producing bacteria depend on the serine palmitoyl transferase (SPT) enzyme, which catalyzes the initial step in the *de novo* synthesis of ceramides, for sphingolipid production. Indeed, the SPT gene is conserved between eukaryotes and prokaryotes and its presence in bacterial genomes has been used as an indication of sphingolipid production. While sphingolipid production is ubiquitous in eukaryotes, it is thought to be restricted to few bacterial phyla. Known sphingolipid-producing bacteria include the Bacteroidetes and Chlorobi phylum, and a subset of Alpha- and Delta-Proteobacteria ^33^. More recently, two additional key enzymes required for bacterial ceramide synthesis have been identified, bacterial ceramide synthase and ceramide reductase ^34^. Phylogenetic analysis of the three bacterial ceramide synthetic genes has identified a wider range of Gram-negative bacteria, as well as several Gram-positive Actinobacteria with the potential to produce sphingolipids ^34^. However, our finding that *P. fluorescens* MYb115, which lacks the SPT gene, produces sphingolipids by the PKS/AMT SgaAB, indicates that there are non-canonical ways of producing sphingolipids in bacteria. Moreover, our analysis of the distribution of the MYb115 PKS SgaAB in bacteria revealed that homologous putative PKS are present in bacteria that are so far unknown sphingolipid producers. This finding strongly suggests that PKS-dependent biosynthesis of sphingolipids is prevalent across bacteria and may even be more broadly distributed than classical SPT-dependent biosynthesis.

By comparing the *C. elegans* transcriptome response to MYb115 and the MYb115 PKS mutant in a metabolic network analysis, we observed an effect of MYb115-derived sphingolipids on host fatty acid and sphingolipid metabolism. Our *C. elegans* lipidomic profiling corroborated the transcriptomic data, providing evidence that MYb115-derived sphingolipids alter *C. elegans* sphingolipid metabolism, resulting in the reduction of certain complex sphingolipid species. A similar effect of gut microbiota-derived sphingolipids on host lipid metabolism was previously observed in mice: *Bacteroides thetaiotaomicron*-derived sphingolipids reduce *de novo* sphingolipid production and increase ceramide levels in the liver ^25^. Also, *B. thetaiotaomicron*-derived sphingolipids alter host fatty acid and sphingolipid metabolism and ameliorate hepatic lipid accumulation in a mouse model of hepatic steatosis ^35^. In humans, bacterial sphingolipid production correlates with decreased host-produced sphingolipid abundance in the intestine and is critical for maintaining intestinal homeostasis ^36^. Thus, interference with host sphingolipid metabolism may be a general effect of bacterial-derived sphingolipids.

What role do MYb115-derived sphingolipids play in host protection against Bt? The current study reveals that MYb115 affects host fatty acid and sphingolipid metabolism. In a previous study we described an association between modulations in fatty acid and sphingolipid metabolism and increased tolerance to Bt infection ^37^. In line with this, we here demonstrate that modulations in sphingolipid metabolism strongly affect survival of infected animals. Our functional genetic analysis of *C. elegans* sphingolipid metabolism enzymes shows that inhibition of *de novo* synthesis of ceramide and inhibition of the conversion of ceramide to glucosylceramides or ceramide-1-phosphate increases survival of *C. elegans* infected with Bt247, while inhibition of the conversion of ceramide to sphingomyelin or sphingosine decreases survival of Bt247-infected animals. Also, MYb115 interacts with host sphingolipid metabolism at least at the conversion of ceramide to sphingomyelin and sphingosine, since the susceptibility phenotypes of the respective mutants are ameliorated or even abrogated in MYb115-treated animals, respectively. Together, these findings provide evidence that MYb115-derived sphingolipids affect *C. elegans* tolerance to Bt247 infection by altering host sphingolipid metabolism. Given that sphingolipids are not only required for the integrity of cellular membranes, but can also act as bioactive signaling molecules involved in regulation of a myriad of cell activities ^38^, their exact roles in microbiota-mediated protection against Bt infection remains to be further explored. Notably, in *C. elegans*, glucosylceramide deficiency was linked to an increase in autophagy ^39,40^, which plays an important role in cellular defence after attack by certain Bt pore-forming toxins (PFTs) ^41^. Also, glucosylceramides serve as a source for the synthesis of complex glycosphingolipids. In *C. elegans*, the Bt-toxin resistant (BRE) proteins BRE-2, BRE-3, BRE-4, and BRE-5 are required for further glucosylation of glucosylceramide, leading to complex glycosphingolipids that are receptors of the *B. thuringiensis* Cry toxin Cry5B ^42^. However, Bt247 only expresses the unique Cry toxin Cry6Ba ^43^, which belongs to the Cry6 family of PFTs ^44^. These proteins are unrelated to Cry5B at the level of their primary sequences and structure ^45^. Also, we could previously exclude an involvement of the *bre* genes in

*C. elegans* defence against Bt247, given that *bre* mutants are susceptible to Bt247 infection ^37^. Still, MYb115-mediated interference with sphingolipid metabolism might affect membrane organisation and dynamics, as well as vesicular transport, which in turn might affect other membrane-associated Bt toxin receptors through modifying their localisation in the plasma membrane. *C. elegans* is thus an ideal experimental system to study the downstream impact of microbiota-derived sphingolipids in the context of pathogen protection, an area that is still largely unexplored ^46^.

## Methods

### *C. elegans* strains and growth conditions

The wildtype *C. elegans* strain N2 (Bristol) ^47^ and all sphingolipid mutant strains were purchased as indicated in Table 1. Worms were grown and maintained on nematode growth medium (NGM) seeded with the *Escherichia coli* strain OP50 at 20 °C, according to the routine maintenance protocol ^48^. Worm populations were synchronised and incubated at 20 °C.

**Table 1.** Worm strains used in this study.

| Worm strain | Genotype | Origin |
| --- | --- | --- |
| N2 |  | CGC |
| RB1036 | *hyl-1(ok976)* | CGC |
| RB1498 | *hyl-2(ok1766)* | CGC |
| RB1487 | *asm-3(ok1744)* | CGC |
| RB1465 | *sptl-1(ok1693)* | CGC |
| RB1579 | *sptl-3(ok1927)* | CGC |
| RB1854 | *sms-1(ok2399)* | CGC |
| FX02613 | *sms-2(tm2613)* | NBRP Tokyo Japan |
| RB2549 | *sms-3(ok3540)* | CGC |
| PHX6977 | *W02F12.2(syb6977)* | SunyBioTech |
| FX00495 | *asah-1(tm495)* | NBRP Tokyo Japan |
| FX00609 | *asah-2(tm609)* | NBRP Tokyo Japan |
| RB1203 | *cerk-1(ok1252)* | CGC |
| VC693 | *cgt-1(ok1045)* | CGC |

### Bacterial strain and growth conditions

The standard laboratory food source *E. coli* OP50 was previously obtained from the CGC. The natural microbiota isolate *Pseudomonas fluorescens* MYb115 (NCBI Reference Sequence: NZ_CP078138.1) isolated from the natural *C. elegans* strain MY379 was used ^16^.

The promoter-exchange strain MYb115 P*_BAD_sga* for targeted in-/activation of the *sgaAB* biosynthetic gene cluster (BGC) was generated *via* insertion of the inducible P*_BAD_* promoter in front of the BGC following an established protocol ^20^. The resulting plasmid (pCEP_kan_*sgaA*) was transformed into the conjugation host *E. coli* ST18 *via* electroporation and introduced into MYb115 *via* conjugation ^49^. The promoter was induced by adding 0.02% (w/v) arabinose (ara) to the culture medium and repressed by adding 0.05% glucose (glc) to the growth medium. Deletions of the single genes *sgaA* and *sgaB* as well as a complete deletion of the whole BGC were carried out following a previously established protocol based on conjugation and homologous recombination ^49,50^. Briefly, fragments upstream and downstream of the target gene were amplified by PCR and assembled into a plasmid using the pEB17 vector ^51^. The resulting plasmids (pEB17_kan_Δ*sgaA,* pEB17_kan_Δ*sgaB* and pEB17_kan_Δ*sgaAB)* were subsequently transformed into the conjugation host *E. coli via* electroporation and the plasmid was introduced into MYb115 *via* conjugation.^49^.

All primer sequences can be found in Table S1. All bacteria were grown on Tryptic Soy Agar (TSA) plates at 25 °C and liquid bacterial cultures were grown in Tryptic Soy Broth (TSB) in a shaking-incubator overnight at 28 °C.

For infection assays with *Bacillus thuringiensis,* we used the strain MYBt18247 (Bt247, our lab strain) and Bt407 (provided by Christina Nielsen-LeRoux, INRA, France) as non-pathogenic control ^37,52^. Spore aliquots of both strains were obtained following a previously established protocol ^53^ with minor modifications ^18^.

### Transcriptome analysis using RNA-seq

Roughly 500 synchronised N2 worms were raised on PFM plates inoculated with MYb115 or MYb115 Δ*sgaAB* (OD_600nm_ of 10) from L1 to L4 stage. At L4 stage worms were transferred to control plates or infection plates (microbiota mixed with Bt247 spores 1:100). Transcriptomic response was assessed 24 h post-transfer, with three independent replicates. Worms were washed off the plates with M9-T (M9 buffer + 0.02% Triton X-100), followed by three gravity washing steps. The worm pellets were resuspended in 800 µL TRIzol (Thermo Fisher Scientific, Waltham, MA United States). Worms were broken up prior to RNA extraction by treating the samples with four rounds of freeze-thaw cycles using liquid nitrogen and a thermo block at 46 °C. The RNA was extracted using Direct-zol™ RNA MicrolPrep (Zymo Research, R2062) and stored at -80 °C.

The RNA was processed by Lexogen (Vienna, Austria) using the 3’ mRNAseq library prep kit and sequenced on an Illumina NextSeq2000 on a P3 flow cell in SR100 read mode. FASTQ files were checked for their quality with MultiQC ^54^, filtered and trimmed with cutadapt ^55^, and aligned to the *C. elegans* reference genome WBcel235 with the STAR aligner (Spliced Transcripts Alignment to a Reference ^56^) followed by an assessment using RseQC ^57^. Ultimately, HTseq-count v0.6.0 ^58^ generated the raw gene counts. The count normalization with the median of ratios method for sequencing depth and RNA composition as well as the analysis for differential expression by a generalized linear model (GLM) was performed using DESeq2 ^59^. Raw data and processed data have been deposited in NCBI’s Gene Expression Omnibus ^60^ and are accessible through GEO Series accession number GSE245296.

### Liquid chromatography-mass spectrometry (LC-MS) analysis of MYb115

For LC-MS analysis, 1 mL liquid culture was harvested *via* centrifugation (1 min, 20 °C, 17,000 x *g*). The cell pellet was resuspended in 1 mL MeOH and incubated at 30 °C for 30 min. The resulting extract was separated from the cell debris *via* centrifugation (30 min, 20 °C, 17,000 x *g*), diluted and submitted to LC-MS measurements. LC-MS measurements were performed on a Dionex Ultimate 3000 (Thermo Fisher Scientific) coupled to an Impact II qToF mass spectrometer (Bruker Daltonics). 5 µL sample were injected and a multistep gradient from 5 to 95% acetonitrile (ACN) with 0.1% formic acid in water with 0.1% formic acid over 16 min with a flow rate of 0.4 mL/min was run (0-2 min 5% ACN; 2-14 min 5-95% ACN; 14-15 min 95% ACN; 15-16 min 5% ACN) on a Acquity UPLC BEH C18 1.7 µm column (Waters). MS data acquisition took place between minutes 1.5 and 15 of the multistep LC gradient. The mass spectrometer was set to positive polarity mode with a capillary voltage of 2.5 kV and a nitrogen flow rate of 8 L/min. We compared the MS^2^ data of compounds **1**-**3** to the MS^2^ data obtained from commercially available sphinganines (sphinganine (d18:0) and sphinganine (d20:0), Avanti Polar Lipids).

### Labeling experiments

Bacterial cultures producing the sphinganine compounds were grown in ISOGRO®-^13^C and ISOGRO®-^15^N (Sigma Aldrich) medium and subsequently analysed by LC-MS to determine the number of carbon and nitrogen atoms, respectively. To confirm the incorporation of serine into the sphinganines, MYb115 P*_BAD_sga* cultures were grown in XPP medium ^51^ with addition of all proteinogenic amino acids (Carl Roth GmbH + Co. KG, Karlsruhe) except serine. To test the incorporation, either ^13^C_3_^15^N-labeled (Sigma Aldrich) serine or regular serine (Carl Roth GmbH + Co. KG, Karlsruhe) displaying the usual isotopic abundances were used. This should result in the production of two isotopologues of each sphinganine. With addition of ^13^C_3_^15^N-labeled serine, the isotopologue that is m_monoisotopic_+3 should be labeled with two ^13^C isotopes and one ^15^N isotope, since one carbon atom is lost through the elimination of CO_2_ during the condensation. In the cultures with regular serine, the isotopologue that is m_monoisotopic_+3 should be labeled with three ^13^C isotopes because of the higher natural abundance of ^13^C compared (1.1%) to ^15^N (0.4%). The two isotopologues, ^13^C_3_ and ^13^C_2_^15^N, were distinguished by their respective masses.

### Metabolic Modeling

For the metabolic model analysis, transcriptomic data was integrated into the iCEL1314 *C. elegans* metabolic model using the MERGE pipeline ^27^ in MATLAB (version: 9.11.0.1769968 (R2021b)) using the COBRA toolbox ^61^). Gene categorization was performed in Python ^62^ (version 3.10.6) using 0.7816 (mu1), 4.856 (mu2), and 8.15 (mu3), as rare, low, and high expression category cutoffs, respectively. Differences between generated metabolic models were assessed by fitting a linear regression model (data ∼ treatment) using Flux Variability Analysis (FVA) ^29^ output (lower bound/upper bound, range per reaction) and Optimal Flux Distribution (OFD) values (equivalent to parsimonious FBA solution) from each model. Significant reactions (alpha = 0.01) from the different data types were combined, and Flux Enrichment Analysis (FEA) was performed to identify significantly affected metabolic model subsystems. For sphingolipid metabolism pathway analysis, FVA was performed on all 23 reactions, with biomass objective minimum set to 50%. Upper bound values were grouped by pathway, then normalized against the mean on the MYb115 flux values for each reaction. Lower bound values were not analysed due to the unidirectional nature of most reactions (lb = 0).

### *P. fluorescens* MYb115 lipidomics

For the bacterial lipidomics experiment, we adapted the extraction method from Brown *et al.* ^36^. 5 mL liquid cultures were incubated for 24 h at 30 °C. The equivalent of 1mL OD_600nm_ of 5 was harvested by centrifugation (1 min, 20 °C, 17,000 x *g*). The cell pellet was resuspended in 0.4 mL H_2_O. 1.5 mL CHCL_3_/MeOH (1:2) were added and the extracts were mixed by vortexing. The cell mixture was incubated at 30 °C with gentle shaking, after 18 h 1mL CHCl_3_/H_2_O (1:1) was added. After phase separation, the organic phase was dried using a nitrogen evaporator and stored at -20 °C.

The relative quantification and annotation of lipids was performed by using HRES-LC-MS/MS. The chromatographic separation was performed using a Acquity Premier CSH C18 column (2.1 × 100 mm, 1.7 μm particle size, VanGuard) a constant flow rate of 0.3 mL/min with mobile phase A being 10 mM ammonium formate in 6:4 ACN:water and phase B being 9:1 IPA:ACN (Honeywell, Morristown, New Jersey, USA) at 40° C. For the measurement, a Thermo Scientific ID-X Orbitrap mass spectrometer was used. Ionization was performed using a high temperature electrospray ion source at a static spray voltage of 3500 V (positive) and a static spray voltage of 2800 V (negative), sheath gas at 50 (Arb), auxiliary gas at 10 (Arb), and ion transfer tube and vaporizer at 325 and 300 °C, respectively.

Data dependent MS^2^ measurements were conducted applying an orbitrap mass resolution of 120 000 using quadrupole isolation in a mass range of 200 – 2000 and combining it with a high energy collision dissociation (HCD). HCD was performed on the ten most abundant ions per scan with a relative collision energy of 25%. Fragments were detected using the orbitrap mass analyser at a predefined mass resolution of 15 000. Dynamic exclusion with an exclusion duration of 5 seconds after 1 scan with a mass tolerance of 10 ppm was used to increase coverage. For lipid annotation, a semi-quantitative comparison of lipid abundance and annotated peaks were integrated using Compound Discoverer 3.3 (Thermo Scientific). The data were normalized to the maximum peak area sum of all samples, the *p*-value per group ratio calculated by a one-way ANOVA with Tukey as post-hoc test, and the *p*-value adjusted using Benjamini-Hochberg correction for the false-discovery rate ^63^. The *p*-values were estimated by using the log-10 areas. The normalized peaks were extracted and plotted using R (4.1.2) within RStudio using the following packages: ggplot2 (3.4.0), readxl (1.4.1), grid (4.1.2), gridExtra (2.3), and RColorBrewer (1.1-3). Metabolomics data have been deposited to the EMBL-EBI MetaboLights database (DOI: 10.1093/nar/gkz1019, PMID:31691833) with the identifier MTBLS8694.

### PKS distribution analysis

The monomodular PKS (KW062_RS19805) and the aminotransferase (KW062_RS19800) in *P. fluorescens* MYb115 (NZ_CP078138) were searched against the non-redundant (nr) National Center for Biotechnology Information (NCBI) database using cblaster (1.8.1) ^23^. PKS encoded by various bacterial genera were aligned and visualised using clinker ^24^.

### C. elegans lipidomics

For lipidomic profiling, N2 worms exposed to MYb115 or MYb115 Δ*sgaAB* were used. Approximately 10,000 worms were raised on either of the bacteria for 70 h until they were young adults. Excess bacteria were removed by three gravity washing steps using M9 buffer. The buffer was thoroughly removed, and the samples were snap-frozen in liquid nitrogen.

Extraction and analysis of lipids were performed as described previously ^64^. Worm pellets were suspended in MeOH and homogenized in a Precellys Bead Beating system (Bertin Technologies, Montigny-le-Bretonneux, France), followed by addition of MTBE. After incubation water was added and through centrifugation the organic phase was collected. The aqueous phase was re-extracted using MTBE/MeOH/H_2_O (10/3/2.5 v/v/v). Organic phases were combined and evaporated to dryness using a SpeedVac Savant centrifugal evaporator (Thermo Scientific, Dreieich, Germany). Proteins were extracted from the residue debris pellets and quantified using a BCA kit (Sigma-Aldrich, Taufkirchen, Germany). Lipid profiling was performed using a Sciex ExionLC AD coupled to a Sciex ZenoTOF 7600 under control of Sciex OS 3.0 (Sciex, Darmstadt, Germany). Separation was achieved on Waters Cortecs C18 column (2.1 mm x 150 mm, 1.6 µm particle size) (Waters, Eschborn, Germany). 40% H_2_O / 60% ACN + 10 mM ammonium formate / 0.1% formic acid and 10% ACN / 90% iPrOH + 10 mM ammonium formate / 0.1% formic acid were used as eluents A and B. Separation was carried out at 40 °C at a flow rate of 0.25 mL/min using a linear gradient as followed: 32/68 at 0.0 min, 32/68 at 1.5 min, 3/97 at 21 min, 3/97 at 25 min, 32/68 at 25.1 min, 32/68 at 30 min. Analysis was performed in positive ionization mode.

Dried samples were re-dissolved in H_2_O/ACN/iPrOH (5/35/60, v/v/v) according to their protein content to normalize for differences in biomass. 10 µL of each sample were pooled into a QC sample. The remaining sample was transferred to an autosampler vial. The autosampler temperature was set to 5 °C and 5 µL were injected for analysis. MS^1^ ions in the *m/z* range 70 to 1500 were accumulated for 0.1 s and information dependent acquisition of MS^2^ was used with a maximum number of 6 candidate ions and a collision energy of 35 eV with a spread of 15 eV. Accumulation time for MS^2^ was set to 0.025 s yielding a total cycle time of 0.299 s. ZenoTrapping was enabled with a value of 80000. QC samples were used for conditioning of the column and were also injected every 5 samples. Automatic calibration of the MS in MS^1^ and MS^2^ mode was performed every 5 injections using the ESI positive Calibration Solution for the Sciex X500 system or the ESI negative Calibration Solution for the Sciex X500 system (Sciex, Darmstadt, Germany).

Data analysis was performed in a targeted fashion for sphingolipids (Table S6). Sphingolipids were identified by manual interpretation of fragmentation spectra following established fragmentation for different sphingolipid classes: *m/z* 268.263491, 250.252926 and 238.252926 for C17iso sphingosine and *m/z* 270.279141, 252.268577 and 288.289706 for C17iso sphinganine based derived sphingolipids. Data analysis was performed in Sciex OS 3.0.0.3339 (Sciex, Darmstadt, Germany). Peaks for all lipids indicated below were integrated with a XIC width of 0.02 Da and a gaussian smooth width of 3 points using the MQ4 peak picking algorithm. All further processing was performed in R 4.2.1 within RStudio using the following packages: tidyverse (v1.3.2), readxl (1.4.1), ggsignif (0.6.4), ggplot2 (3.3.6), scales (1.2.1). Significance was tested using a Welch-Test within ggsignif. Metabolomics data have been deposited to the EMBL-EBI MetaboLights database (DOI: 10.1093/nar/gkz1019, PMID:31691833) with the identifier MTBLS8440.

### Bt survival assay

*B. thuringiensis* survival assays were performed as described previously with minor adjustments ^18,65,66^. N2 wildtype worms and the sphingolipid mutants were synchronised and grown on PFM plates seeded with 1 mL MYb115 or OP50 (OD_600nm_ of 10) until they reached the L4 stage. Infection plates were inoculated with each of the bacteria adjusted to OD_600nm_ of 10 mixed with Bt247 spores or Bt407 For the infection L4 worms were washed off the plates with M9 buffer and 30 worms were pipetted onto infection plates and incubated at 20 °C. To assess survival, all worms were counted as either alive or dead 24 h after infection. Worms were considered dead if they did not respond to light touch with a platinum wire picker. We plotted all survivals as survival curves (Figure S6) but provided a summary of the data in a heatmap (Figure 5C). The area under the survival curve (AUC) was calculated for the *C. elegans* mutant strains and the mean AUC of *C. elegans* wildtype N2. The AUC for the mutant strain was then subtracted from the mean AUC of wildtype worms (ΔAUC). Based on the ΔAUC values, the shading for the heatmap was determined (Figure 5B). Bt survival assays were done each with three to four replicates per treatment group and around 30 worms per replicate for each independent experiment. Statistical analyses were performed with RStudio (Version 4.1.2) ^67^. GLM analysis with Tukey multiple comparison tests ^68^ and Bonferroni ^69^ correction were used for all survival assays individually. For overall survival, represented in the heatmap (Figure 5C) GLM with False Discovery Rate (FDR) ^63^ were used. Graphs were plotted using ggplot2 ^70^ and were edited in Inkscape (Version 1.1).

### Bacterial colonization assay

To test for differences in colonization of *C. elegans* L4 and young adults by MYb115 and MYb115 Δ*sgaAB*, colonization was quantified by counting colony forming units (CFUs). Worms were exposed to MYb115 and MYb115 Δ*sgaAB* from L1 to L4 larval stage or additionally 24 h until worms reached young adulthood. To score the CFU, worms were washed off their plates with M9-T (M9 buffer + 0.025% Triton-X100) followed by five gravity washing steps with M9-T. Prior to soft bleaching, worms in M9-T were paralyzed with equal amounts of M9-T and 10 mM tetramisole to prevent bleach solution entering the intestine. Worms were bleached for two min with a 2% bleach solution (12% NaClO diluted in M9 buffer). Bleaching was stopped by removing the supernatant and washing the samples with PBS-T (PBS: phosphate-buffered saline + 0.025% Triton-X100). A defined number of worms was transferred into a new tube with PBS-T. A subsample of this was used as a supernatant control, while the remaining sample was homogenized with sterile zirconia beads (1 mm) using the BeadRuptor 96 (omni International, Kennesaw Georgia, USA) for 3 min at 30 Hz. Homogenized worms were diluted (1:10/1:100) and plated onto TSA plates, as well as the undiluted supernatant as control. After 48 h at 25 °C, colonies were counted and the CFUs per worm were calculated. To determine significant differences, we performed a *t*-test.

### Pumping behavior

To score the pumping rate, i.e. the back and forth movement of the grinder, worms were exposed to either MYb115 or MYb115 Δ*sgaAB*. Pumping was scored at L4 larval stage, young adults and young adults infected with Bt247 (1:100). Only worms that were on the bacterial lawn were counted for a period of 20 s. 15-20 worms per condition were counted. To determine significant differences, we performed pairwise Wilcoxon test.

## Acknowledgements

We thank Lena Bluhm, Laura Brügmann, Johanna Jarstorff, Hanne Griem-Krey and Sabrina Butze for their technical support. This work was funded by the German Science Foundation DFG (Collaborative Research Center CRC1182 Origin and Function of Metaorganisms, project A1.2 to KD, project A1.1 to HS, and project A1.5 to CK). We thank the Caenorhabditis Genetics Center (University of Minnesota, Minneapolis, Minnesota, USA), funded by the NIH Office of Research Infrastructure Programs (P40OD010440) for *C. elegans* strains. Work in the Bode lab was partially supported by an ERC advanced grant (835108) and the Max-Planck Society. Work in the Metabolomics and Proteomics Core and Research Unit Analytical BioGeoChemistry, Helmholtz Zentrum München was partially supported by the German Science Foundation DFG (Project number 431572533 (MetClassNet) to Michael Witting).

## Extended data Figures

**Figure S1.**
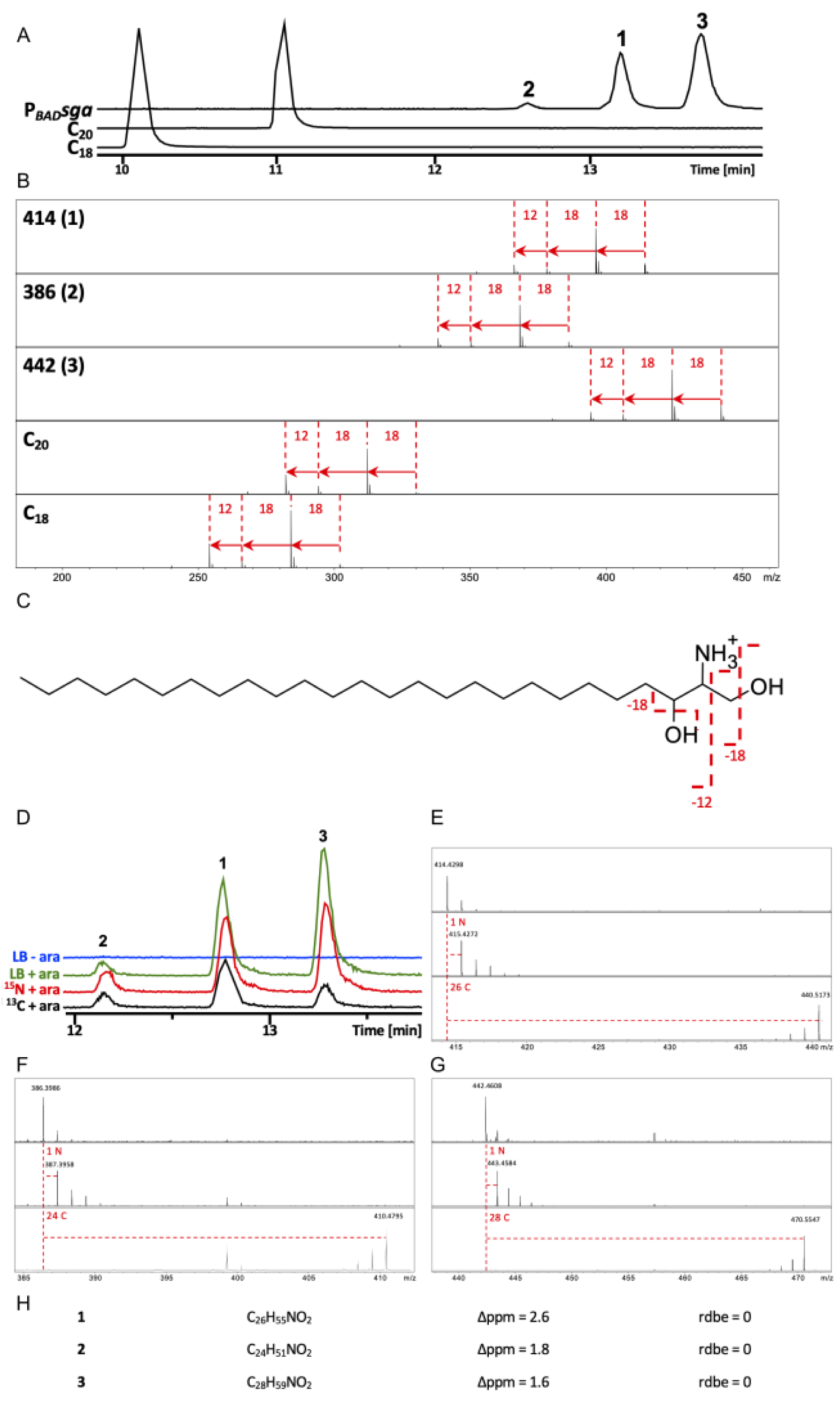
MYb115 PKS SgaAB-derived compounds 1-3 are very long chain sphinganines. **(A)** Extracted ion chromatograms of compounds **1**, **2** and **3**, as well as sphinganine (d18:0) and sphinganine (d20:0). **(B)** Fragmentation patterns of **1**, **2** and **3**, as well as sphinganine (d18:0) and sphinganine (d20:0). **(C)** Fragmentation of sphinganines, exemplary shown for the structure of compound **1**. **(D-H)** Sum formula determination of compounds **1**, **2** and **3** using isotopic labeling and LC-MS. **(D)**: Extracted ion chromatogram (EICs) of compounds **1**, **2** and **3** in unlabeled samples with (green) and without arabinose (blue), as well as ^15^N- (red) and ^13^C-labeled (black) samples. **(E-G)**: Mass shifts compared to LB cultivation (dashed red lines) represent the number of carbon and nitrogen atoms incorporated. **(H)** Sum formula and structural data of compounds **1-3**.

**Figure S2:**
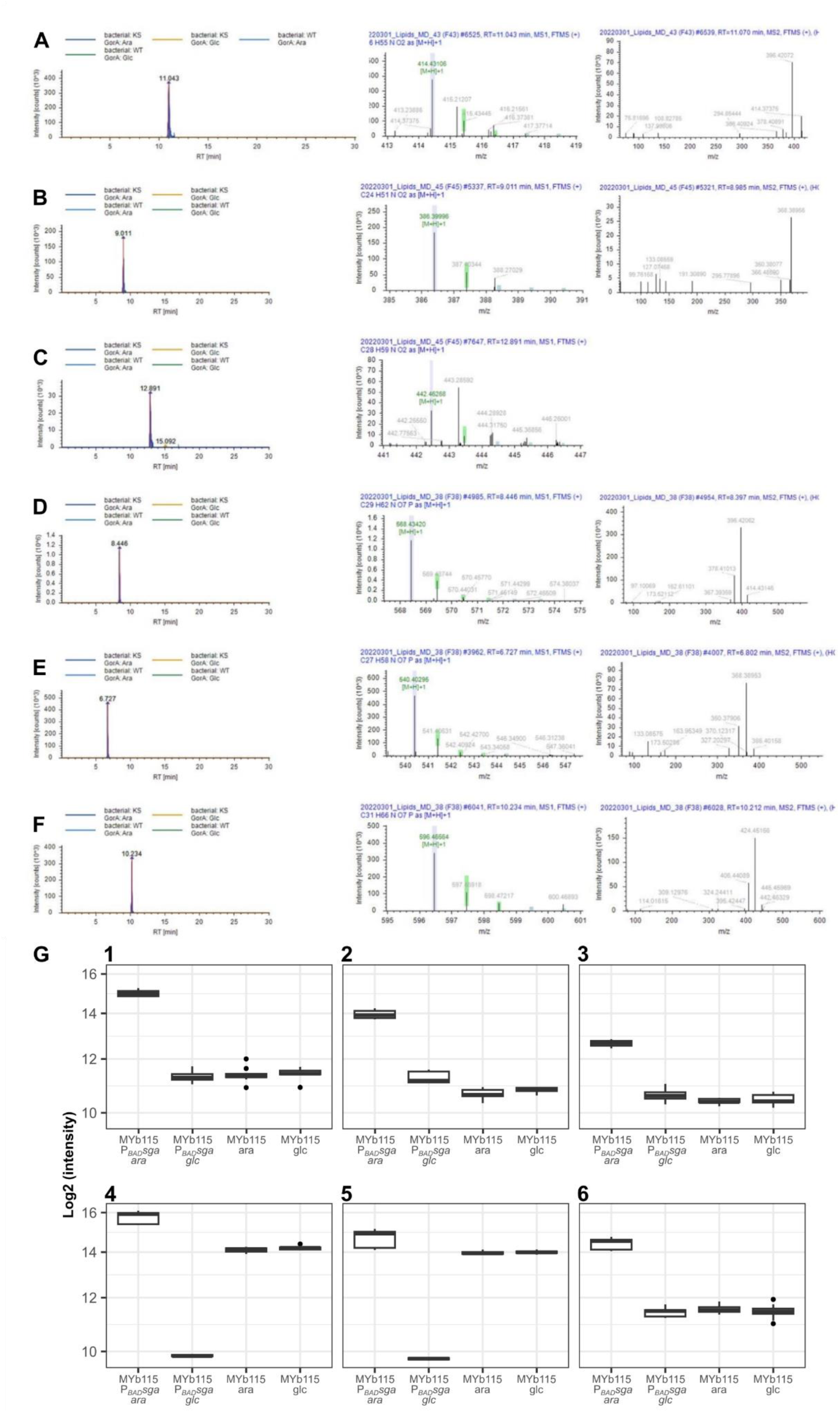
*P. fluorescens* MYb115 PKS produces long chain sphinganines and phosphoglycerol sphingolipids. Original LC-MS data of compounds **1** (**A**), **2** (**B**), **3** (**C**), **4** (**D**), **5** (**E**) and **6** (**F**) stemming from the lipidomics experiments. Left: LC chromatogram; Middle: MS spectra; Right: MS^2^ spectra. Data was extracted using Compound Discoverer 3.3. **(G)** Relative abundance of the sphinganine compounds **1**, **2** and **3** and the phosphoglycerol sphingolipids **4**, **5** and **6** in MYb115 wt and MYb115 P*BADsga* in the presence of arabinose (ara) for activation or in the presence of glucose (glc) for repression of transcription of P*BADsga*.

**Figure S3:**
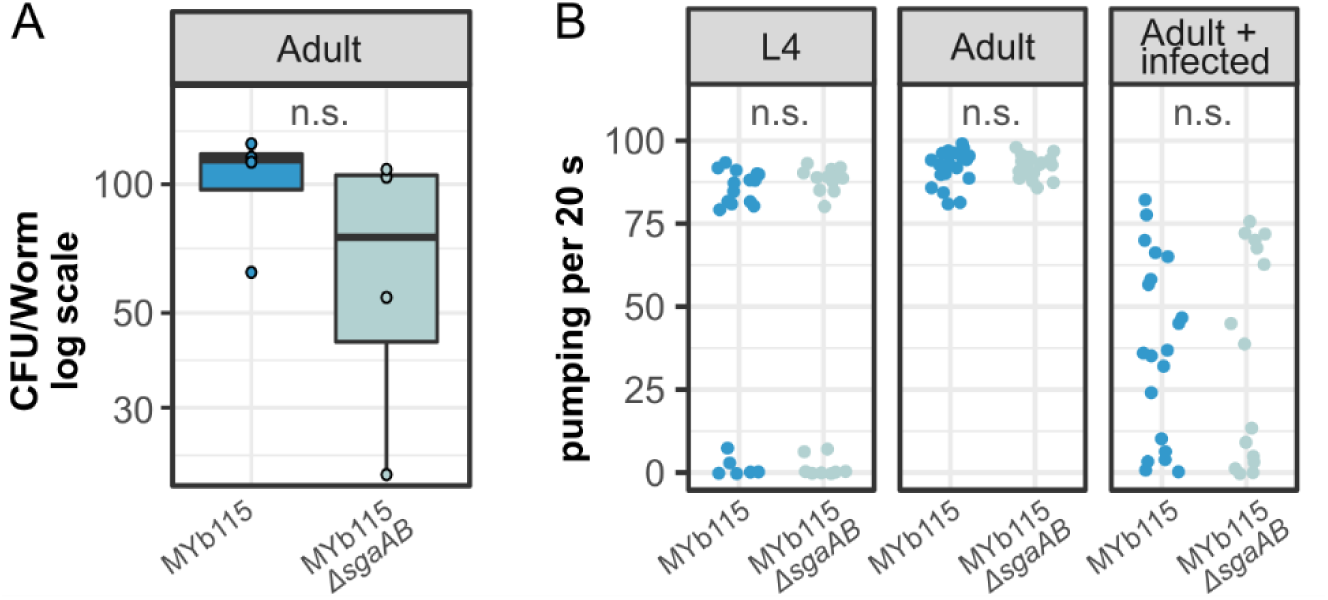
MYb115-derived sphingolipids do not affect host colonization or *C. elegans* feeding behavior. **(A)** Bacterial load of adult worms exposed to either MYb115 or MYb115 Δ*sgaAB*. No significant difference between the colonization of the worm between the two bacterial treatments. *t*-test was performed (*p* = 0.1625). **(B)** Pumping of different worm stages either on MYb115 or MYb115 Δ*sgaAB.* No significant difference between the pumping of worms fed with MYb115 or MYb115 Δ*sgaAB* depending on the given larval stage/ treatment, pairwise Wilcoxon test was performed *(p =* 1.000).

**Figure S4.**
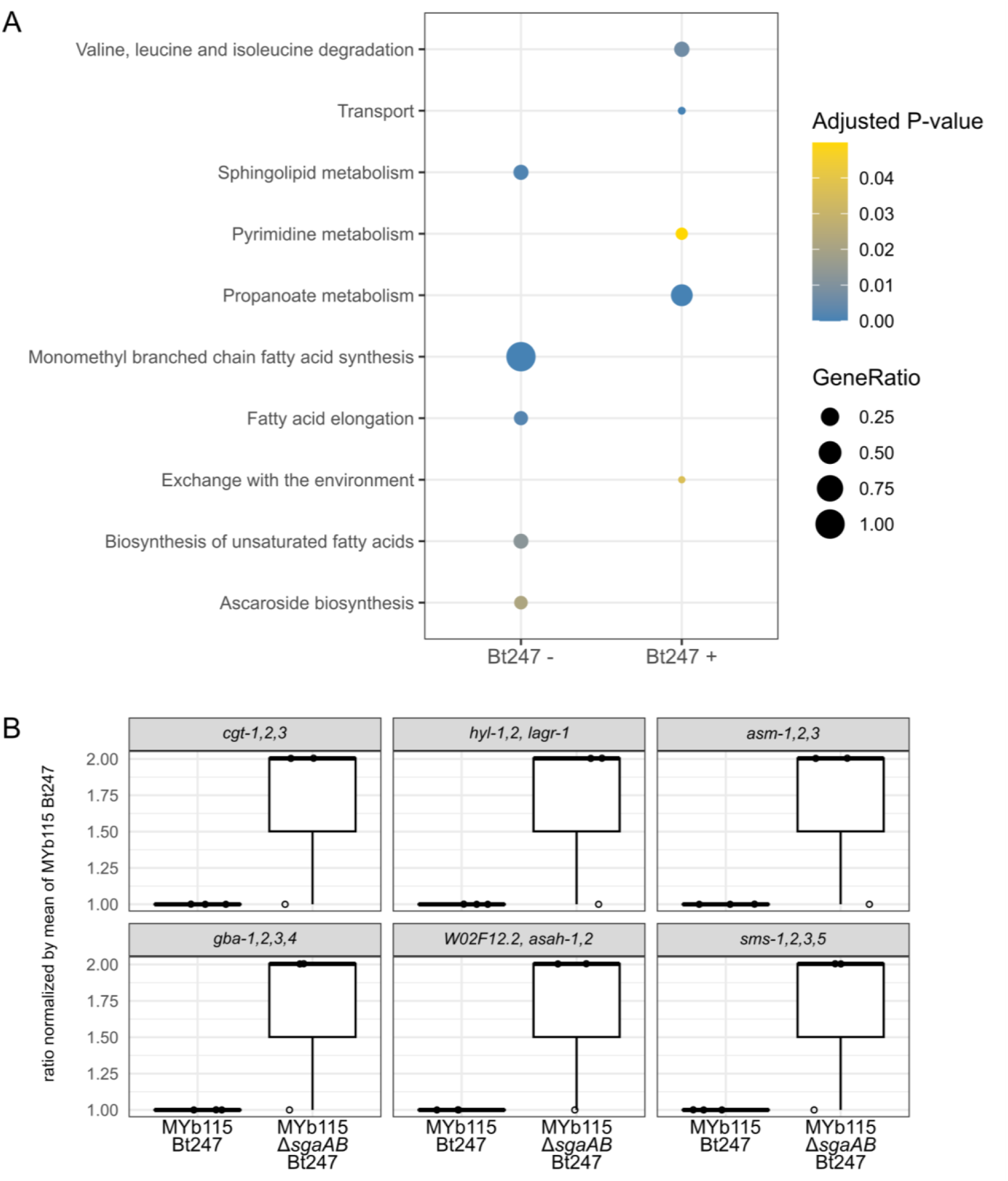
Metabolic network analysis reveals that MYb115-derived sphingolipids affect host fatty acid and sphingolipid metabolism. **(A)** Flux enrichment analysis results in the absence (Bt247-) and presence (Bt247+) of the pathogen. Significant reactions comparing mutant and WT conditions from linear regression models of all 3 data types (ub, lb, OFD) were combined (while removing duplicates) and used against the background of all reactions within the iCEL1314 model. Enrichment was performed with the FEA function in the COBRA toolbox. **(B)** Ratio of upper bound (ub) values for six reactions encoded by the *C. elegans* sphingolipid metabolism enzymes *cgt-1*, *2*, *3*, *hyl-1*, *2* and *lagr-1*, *asm-1*, *2*, *3*, *gba-1*, *2*, *3*, *4*, *W02F12.2* and *asah-1*, *2*, and *sms-1*, *2*, *3*, *5* that all have ceramide as a substrate or product. FVA upper bound values normalized by mean upper bound value of the MYb115_Bt247 group for each reaction.

**Figure S5.**
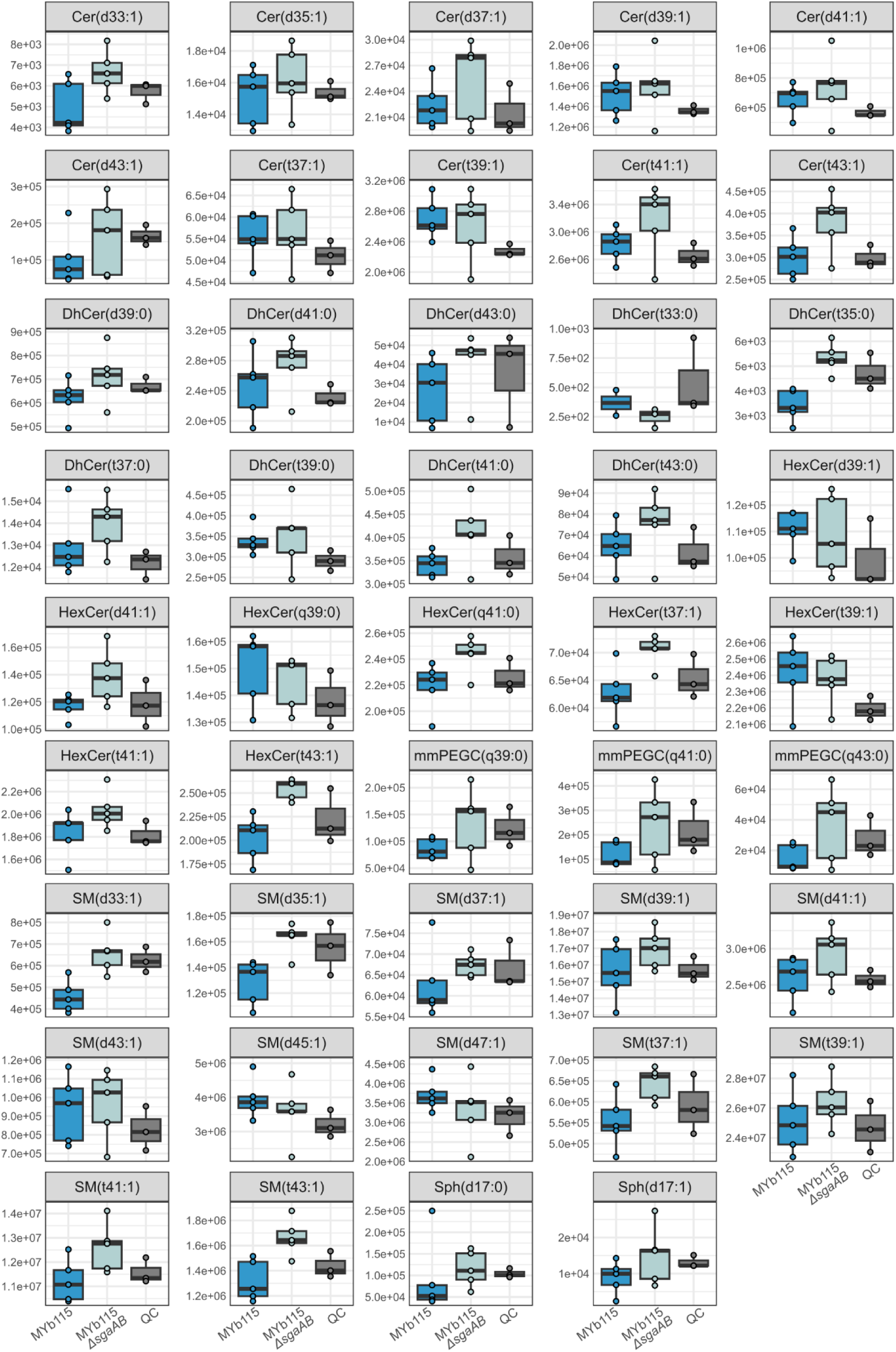
Effects of MYb115-derived sphingolipids on C. elegans sphingolipid profiles. The boxplots show the difference in ratio of different sphingolipids in worms exposed to MYb115 Δ*sgaAB* and MYb115, the data is summarised in the heatmap (**Figure 6C**). Dihydroceramides (DhCer), Ceramides (Cer), Sphingomyelins (SM), Hexosylceramides (HexCer), with hydroxylated fatty acyls (t) or non-hydroxylated fatty acyls (d), Hexosylceramides with phytosphingosine base and hydroxylated fatty acyls (HexCer(q)), monomethyl phosphoethanolamine glucosylceramide (mmPEGC(q)).

**Figure S6:**
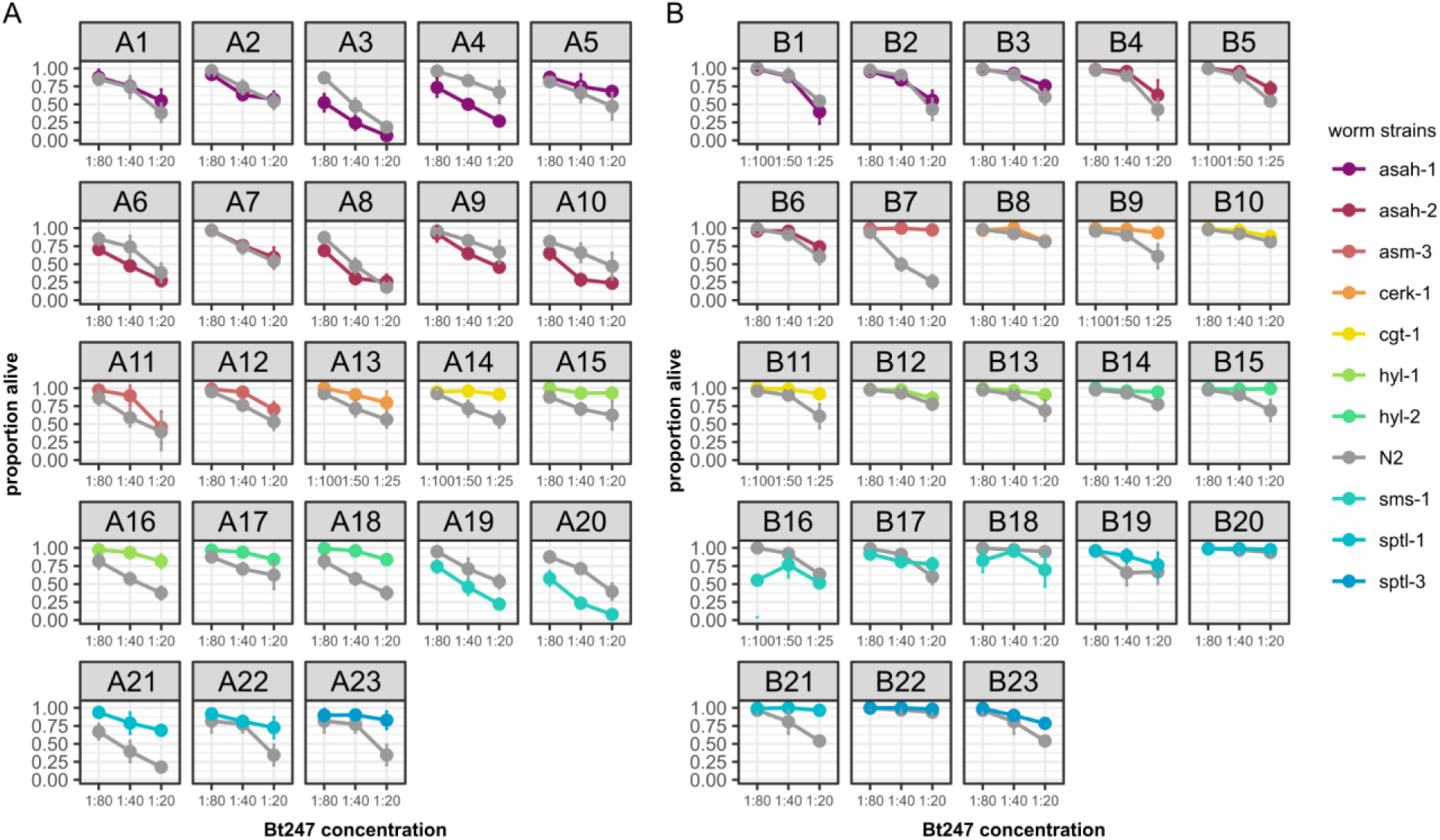
Overview of all individual survival assays from heatmap (Figure 5C), comparing the survival of N2 *versus* the different *C. elegans* sphingolipid metabolism mutants on either OP50 **(A)** or MYb115 **(B)**. Means ± standard deviation (SD) of n = 4, are shown in all survival assays. Statistical analyses were carried out with the GLM framework and Bonferroni adjustment for multiple testing, \*\*\**p* < 0.001. All *p*-values can be found in Table S7.

## Notes

### Competing Interest Statement

The authors have declared no competing interest.

